# Avian keratin disorder of Alaska black-capped chickadees is associated with Poecivirus infection

**DOI:** 10.1101/284992

**Authors:** Maxine Zylberberg, Caroline Van Hemert, Colleen M. Handel, Joseph L. DeRisi

## Abstract

**Background:** Avian keratin disorder (AKD) is an epizootic of debilitating beak deformities, first documented in black-capped chickadees (*Poecile atricapillus*) in Alaska during the late 1990s. Similar deformities have now been recorded in dozens of species of birds across multiple continents. Despite this, the etiology of AKD has remained elusive, making it difficult to assess the impacts of this disease on wild populations. We previously identified an association between infection with a novel picornavirus, Poecivirus, and AKD in a small cohort of black-capped chickadees.

**Methods:** To test if the association between Poecivirus and AKD holds in a lager study population, we used targeted PCR followed by Sanger sequencing to screen 124 symptomatic and asymptomatic black-capped chickadees for Poecivirus infection. We further compared the efficacy of multiple non-terminal field sampling methods (buccal swabs, cloacal swabs, fecal samples, and blood samples) for Poecivirus screening. Finally, we used both *in situ* hybridization and a strand-specific expression assay to localize Poecivirus to beak tissue of AKD-positive individuals and to determine if virus is actively replicating in beak tissue.

**Results:** Poecivirus was detected in 29/29 (100%) individuals with AKD, but only 9/95 (9.5%) asymptomatic individuals with apparently normal beaks (p < 0.0001). We found that cloacal swabs are the most sensitive of these sample types for detecting Poecivirus in birds with AKD, but that buccal swabs should be combined with cloacal swabs in evaluating the infection status of asymptomatic birds. Finally, we used both *in situ* hybridization and a strand-specific expression assay to localize Poecivirus to beak tissue of AKD-positive individuals and to provide evidence of active viral replication.

**Conclusion:** The data presented here show a strong, statistically significant relationship between Poecivirus infection and AKD, and provide evidence that Poecivirus is indeed an avian virus, infecting and actively replicating in beak tissue of AKD-affected BCCH. Taken together, these data corroborate and extend the evidence for a potential causal association between Poecivirus and AKD in the black-capped chickadee. Poecivirus continues to warrant further investigation as a candidate agent of AKD.

## BACKGROUND

In recent years, beak deformities have been documented in dozens of avian species across continents. Birds afflicted by this disease, called avian keratin disorder (AKD), develop beak deformities characterized by elongation and often crossing and marked curvature (Figure 1) [1]. These deformities result in decreased ability to feed and preen, changes in diet, and higher susceptibility to infection with a variety of parasites and pathogens, and ultimately lead to decreased fitness and survival [1-5]. While the population-level impacts of AKD remain uncertain, the high prevalence, fitness impacts, and widespread nature of AKD among multiple host species raise concern that this pathology could have broad-ranging and negative impacts on wild bird populations [1, 6, 7].

**Figure 1.**
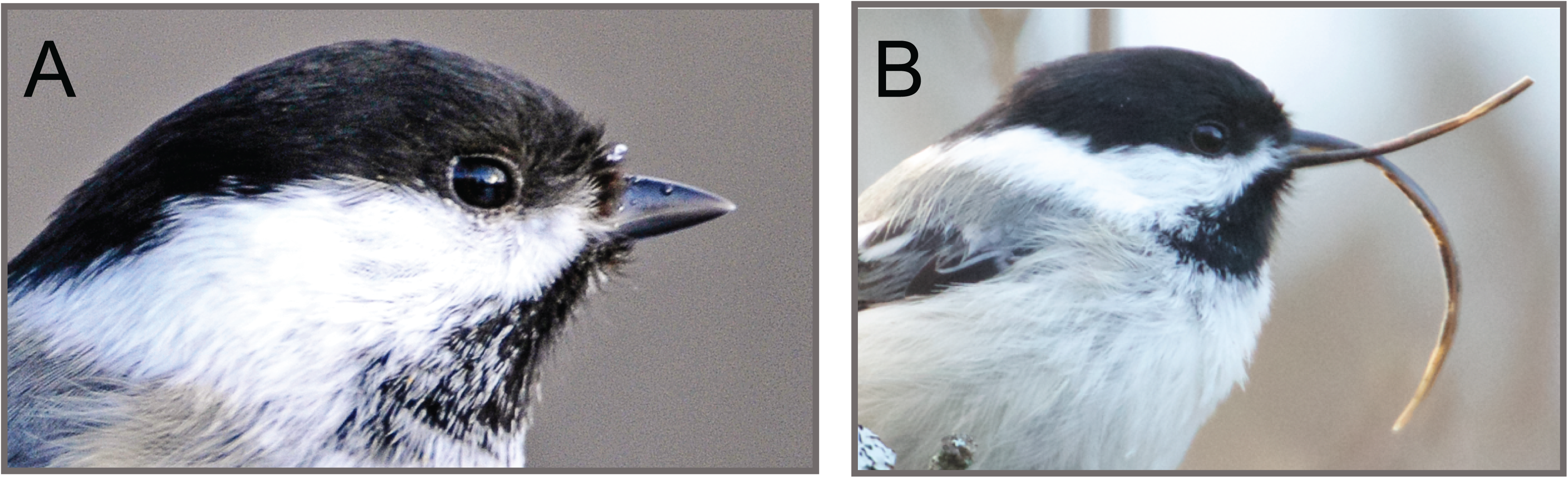
Avian keratin disorder. Panel A) BCCH with a normal beak; photo by John Schoen. Panel B) BCCH exhibiting beak overgrowth characteristic of AKD; photo by Martin Renner.

AKD was first documented among black-capped chickadees (BCCH, *Poecile atricapillus*) in Alaska in the late 1990s [1], with an average prevalence of 6.5% in adult Alaskan BCCH. Meanwhile, morphologically similar deformities have been documented in more than 40 avian species in North America and over 30 species in the United Kingdom [1, 8-10]. Such deformities appear to be particularly common in corvids (such as the northwestern crow [*Corvus caurinus*] in North America and the rook [*C. frugilegus*] in the United Kingdom); cavity-nesting passerines (such as BCCH and the red-breasted nuthatch [*Sitta canadensis*] in North America and the Eurasian blue tit [*Cyanistes caeruleus*] in the United Kingdom); and raptors in North America [1, 6]. Despite the similarity of the gross pathology observed across species, it is unknown if a common factor is responsible. This is partly because the cause of AKD has remained elusive for over two decades [3, 7].

The identification of the causative agent of AKD is necessary to determine whether the beak pathologies observed across species represent a multi-species epizootic. Furthermore, knowing the etiologic agent will allow scientists to determine the prevalence of this disease and evaluate its impact on avian populations apart from other potential causes of gross beak abnormalities. Indeed, a variety of factors can contribute to beak deformities, including environmental contaminants, nutritional deficiencies, trauma, and exposure to infectious agents [11]. However, over the years, multiple studies failed to find clear evidence of contaminant exposure, a nutrient deficiency, or bacterial or fungal infection underlying AKD [3, 12]. In 2016, we used unbiased, high-throughput metagenomic sequencing of beak tissue from BCCH affected by AKD to identify Poecivirus, a candidate agent. Poecivirus is most closely related to avian picornaviruses, but represents a novel viral genus [7]. Subsequent screening of 28 BCCH revealed that 19/19 of BCCH affected by AKD tested positive for Poecivirus, compared with only 2/9 asymptomatic individuals. These results suggested that Poecivirus merited further investigation as a candidate etiological agent of AKD in BCCH.

Our previous work indicated that in addition to beak tissues, Poecivirus could be detected in cloacal and buccal swabs, providing a non-terminal method of sampling individuals for infection, and a potential route to sampling a substantially larger number of individuals [7]. Here, we apply these and other methods to further explore the association of Poecivirus and AKD. We tested a variety of samples (cloacal and buccal swabs, blood, and feces) from 29 individuals affected by AKD and 95 asymptomatic control individuals for the presence of Poecivirus using targeted PCR primers followed by Sanger sequencing. In addition to increasing our sample size for correlation between AKD and Poecivirus, we used quantitative PCR (qPCR) to investigate the relationship between viral load in beak tissue and the extent of beak deformities observed in AKD cases. Finally, we used *in situ* hybridization to localize viral particles and a strand-specific gene expression assay to demonstrate the presence of replicating virus in beak tissue of individuals with AKD. Taken together, the data presented here provide additional evidence to support our hypothesis that Poecivirus is a potential candidate etiological agent of AKD in black-capped chickadees.

## METHODS

### Sample collection

We tested 124 BCCH (29 individuals with AKD and 95 asymptomatic control individuals) for the presence of Poecivirus using cloacal swabs. To obtain samples, individuals were captured using funnel traps and mist nets in Anchorage and the Matanuska-Susitna Valley, Alaska, during the non-breeding season in 2016 (March-April, October-December) and 2017 (March, April). Standard beak measurements were used to classify individuals as AKD-affected or unaffected [1]. Briefly, an individual was considered AKD-affected if it had a nares-to-tip length (chord measurement from anterior end of the right nare to the tip of the upper beak) ≥8.25 mm, an overbite or underbite of >1.0 mm, or, in the case of specimens for which we had beak samples, if its beak exhibited evidence of hyperkeratosis at the cellular level [7]. We chose to use cloacal swabs because they are non-destructive and easily obtained, and we previously showed cloacal swabs to contain relatively high viral load [7]. In addition, we tested buccal swabs, blood samples, and fecal samples from a subset of individuals to compare the efficacy of a variety of non-terminal sample types for Poecivirus detection. We tested buccal swabs from 21 of the individuals with AKD and 75 of the control individuals. We tested blood samples collected by brachial venipuncture and fecal samples obtained opportunistically from subsets of individuals with AKD (N = 13 and N = 4, respectively). Swab, blood, and fecal samples (collected opportunistically if a bird defecated during handling) were placed in Longmire buffer [13] and stored frozen at −80°C until processed.

For viral load determination, *in-situ* hybridization, and gene expression assay, we used samples remaining from our previous study [7], for which we obtained beak tissue samples from 28 individuals. Nineteen of these exhibited AKD and were trapped using funnel traps and mist nets in Anchorage and the Matanuska-Susitna Valley, Alaska, during the non-breeding season from 2001-2015. Data on infection status of these individuals have been published previously. The 10 individuals with AKD that were collected from 2001-2010 were euthanized upon capture with isoflurane using the open-drop method and stored frozen at −20°C; the remaining 9 specimens were captured in the winter of 2014 and spring of 2015, euthanized, and stored overnight at 4°C prior to necropsy, at which time portions of tissues were frozen at −80°C and additional samples were placed in formalin. From 2 of these, we collected cloacal and buccal swabs prior to euthanasia; swabs were stored in Longmire buffer [13]. Nine individuals not affected by AKD were collected opportunistically between 1995 and 2010 and stored frozen at - 20°C; tissues from a subset of these individuals were used as negative controls in the gene expression assay measuring viral replication. Of the tissue samples collected from our previous study, we had adequate samples remaining from fifteen individuals that tested positive for Poecivirus; of these 14 exhibited AKD, and one was asymptomatic.

All work was conducted with the approval of the U. S. Geological Survey (USGS) Alaska Science Center Institutional Animal Care and Use Committee (Assurance #2016-14) and under appropriate state and federal permits.

### Testing for an association between Poecivirus and avian keratin disorder

RNA was extracted from cloacal and buccal swab samples using a Zymo viral RNA kit. For each sample, 75 μl of Longmire buffer was mixed with 225 μl of viral RNA buffer and the extraction proceeded as described in the manufacturer’s protocol. RNA from tissue samples was extracted using a Zymo quick RNA miniprep kit as previously described [7]. Following extraction, 200 ng of RNA were reverse transcribed in 10 μl reactions containing 100 pmol random hexamer, 1 ? reaction buffer, 5 mM dithiothreitol, 1.25 mM (each) deoxynucleoside triphosphates (dNTPs), and 100 U Superscript III (Life Technologies); mixtures were incubated at 25°C for 5 min, 42°C for 60 min, and 70°C for 15 min. Following reverse transcription, cDNA was screened via PCR using Poecivirus-specific primers (Additional File 1). Samples were initially screened via qPCR using primers Poeci_7F and Poeci_7R; samples with a positive result via qPCR were than subjected to PCR using the additional primer pairs to obtain longer amplicons that were Sanger sequenced to confirm the presence of Poecivirus. Quantitative PCR mixtures contained 1X LC480 Sybr Green master mix (Roche), 0.1 μM Poeci_7F, 0.1 μM Poeci_7R, and 5μl of 1:20- diluted cDNA. PCR mixtures contained 1? iProof master mix, 0.5 μM primer, and 5 μl sample template. Thermocycling consisted of 98°C for 30 sec, then 40–45 cycles of 98°C for 10 sec, 58°C for 10 sec, and 72°C for 30 sec, then a 5 min elongation step of 72°C. Amplicons were visualized using agarose gels; those in the correct size range were purified using a DNA Clean and Concentrator kit or Zymoclean Gel DNA recovery kit (Zymo) and Sanger sequenced (Quintara Biosciences). We used likelihood ratio chi-square tests to test whether there was a difference in likelihood of infection with Poecivirus between AKD and control individuals.

### Viral load

Quantitative PCR was conducted as described above and was used to determine viral load in beak samples from Poecivirus-positive individuals. For each sample, the cyclic threshold (Ct) value obtained using primers Poeci_7F and Poeci_7R (Ct_Poeci_) was normalized to the Ct_ref_ value obtained using primers to avian cellular RNA (avi_8F, avi_8R). Viral load was calculated using the equation viral load = 2^(CtPoeci – Ctref)^. We used linear regression to test for a relationship between beak length and viral load in Poecivirus-positive individuals.

### Virus localization of mRNA via *in situ* hybridization

For mRNA *in situ* hybridization, we examined tissue from four of the individuals with the highest qPCR measured levels of Poecivirus in beak tissue. Beak tissue was collected and stored frozen at −80°C until processed; Tissue was fixed overnight in 4% paraformaldehyde at room temperature, and then decalcified for 2.5 weeks in 0.5 mM EDTA at 4°C. Decalcified tissue was frozen in Optimal Cutting Temperature (OCT) embedding media with dry ice and cut into 10 μm sections on a cryostat. *In situ* detection of mRNA was conducted using the RNAscope 2.5 High Definition (HD) Brown assay kit (Advanced Cell Diagnostics, Inc., Hayward, CA). The assay was conducted following the manufacturer’s recommendations, with the exception that sections underwent a 45 min protease plus treatment in place of the recommended 30 min treatment. Images were captured with a Nikon Ti-E microscope fitted with a Nikon DS-Ri2 camera.

### Detection of viral replication

The presence of negative strand virus was evaluated using the NanoString nCounter™ Elements system (NanoString, Seattle, WA) gene expression assay. Eight oligonucleotide probes were designed to target negative strand virus evenly distributed across the viral genome, 2 oligonucleotide probes were designed to target positive strand virus, and an additional 2 oligonucleotide probes were designed to target a BCCH housekeeping gene, NADH ND2 (GenBank: KF183899.1); oligonucleotide probes were obtained from IDT (Coralville, IA). Negative and positive strand virus genome segments were made using Gibson assembly cloning followed by transcription with T7 polymerase [14]; equimolar pools of this RNA served as positive controls of probe activity. A 50 ng sample was hybridized with Tagsets (NanoString) and oligonucleotide probes at 65°C for 16 hours and the excess probes were washed away; in addition, a water negative control was run in one lane, and positive control RNA was run in each of four lanes, with 100, 1,000, 10,000, or 100,000 copies of target RNA added to control lanes 1-4, respectively. The digital counts were captured by nCounter Digital Analyzer and the data were analyzed by nSolver Analysis Software (NanoString). Following standard NanoString analysis methods, individuals were considered to have a positive result for any single probe if that probe exhibited levels 2 standard deviations or greater above that of negative controls. We used linear regression to test for a relationship between beak length and viral RNA count in Poecivirus-positive individuals. Statistical analyses were conducted using JMP 7.0.1 by SAS (SAS Institute Inc., Cary, NC).

## RESULTS

Among the BCCH with AKD, 29/29 (100%) individuals tested positive for Poecivirus by PCR of cDNA from buccal or cloacal swab samples or both. Only 9/95 (9.5%) control individuals tested positive for the presence of Poecivirus (Figure 2). Of note, follow-up data are available for four of the asymptomatic control individuals that tested positive for Poecivirus. One of these (1/4, 25%) had developed an elongated beak typical of AKD when recaptured 6 months later (beak length of 12.5 mm). The other three birds (3/4, 75%) did not show evidence of elongated beaks when recaptured 6-12 months after testing positive for Poecivirus; their maximum subsequent beak lengths were 7.8, 7.4, and 7.1 mm (we do not have subsequent data on Poecivirus infection status).

**Figure 2.**
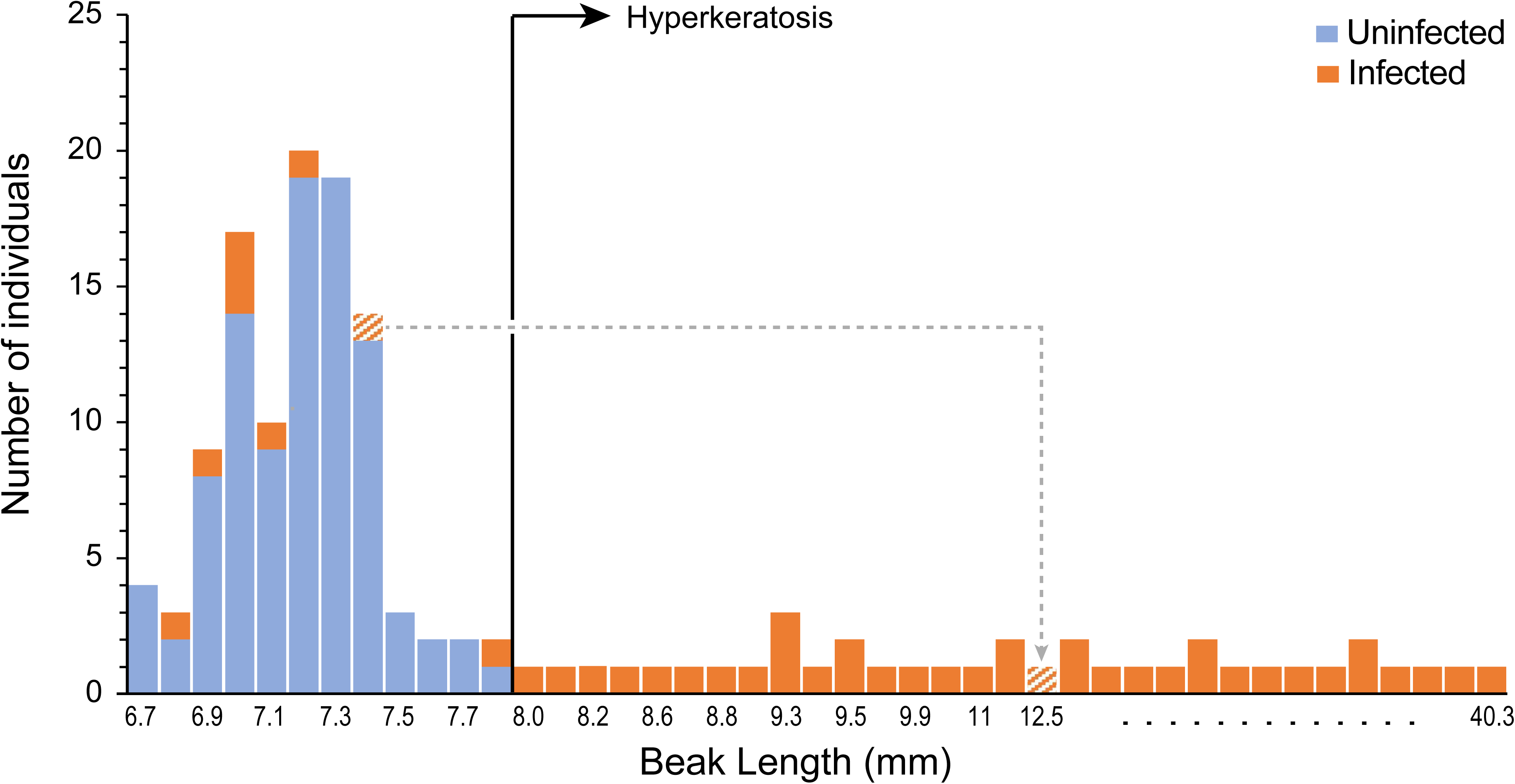
Association between Poecivirus and AKD in BCCH. Y-axis shows the number of individuals tested with a given beak length; individuals testing negative for Poecivirus are highlighted in blue and those testing positive for Poecivirus are in orange. Note that the x-axis is not to scale (beak lengths for which we had no individual data are excluded from the graph). Individuals to the right of the vertical line are classified as having AKD based on beak length or morphology. The hatched data point represents the single individual with an apparently normal beak that tested positive for Poecivirus and later developed an elongated beak; this is the only individual represented twice on the graph and the data points are linked by a dashed line, with the arrow pointing to the later beak measurement.

Among the BCCH with AKD, 28/29 (96.6%) tested positive via cloacal swab. In comparison, 3/29 (10.4%) individuals with AKD tested positive for Poecivirus via buccal swab, one of these having tested negative via cloacal swab (Table 1). Among the asymptomatic control BCCH, 5/95 (5.3%) individuals tested positive for Poecivirus via cloacal swab, 4/75 (5.3%) tested positive via buccal swab, and none tested positive via both cloacal and buccal swab. None of the blood samples (0/13) or fecal samples (0/4) from AKD-positive individuals tested positive for Poecivirus, despite 13/13 of these individuals testing positive for the presence of Poecivirus via cloacal and/or buccal swab. Overall, targeted PCR testing of cloacal swabs for the presence of Poecivirus resulted in most cases of Poecivirus detection, and were most consistent with our previous, beak tissue-based testing, which detected the presence of Poecivirus in 19/19 (100%) of AKD individuals tested [7]. However, cloacal and buccal swabs provided complementary information on the presence of Poecivirus in individuals without apparent beak deformities.

**Table 1:**
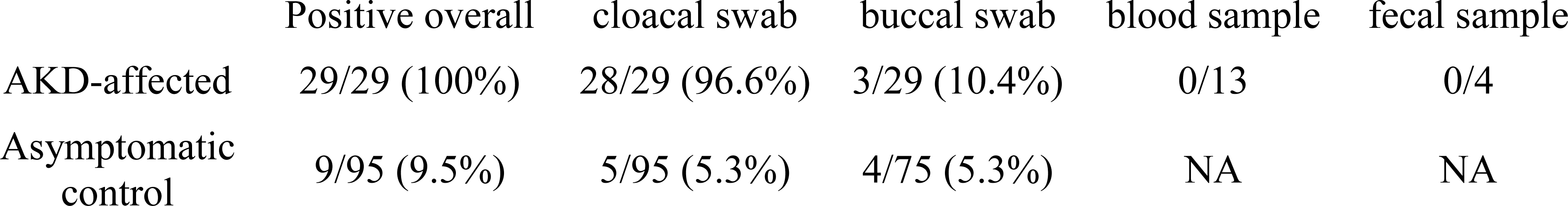
Efficacy of different non-terminal sampling methods for detecting Poecivirus in black-capped chickadees (*Poecile atricapillus*) exhibiting signs of avian keratin disorder (AKD-affected) and in asymptomatic controls. NA = not applicable.

Individuals with AKD were significantly more likely to be infected with Poecivirus than expected by chance if we consider the swab data from the current study (p < 0.0001, χ^2^ = 93.29, df = 1, N = 124) and when we combine the swab data from the current study with our previous data that tested individuals for Poecivirus using beak and cloacal tissue (p < 0.0001, χ^2^ = 129.03, df = 1, N = 150; Figure 2) [7]. However, we found only a weak, if statistically significant, relationship between beak length and viral load in beak tissue (p = 0.026, R^2^_adj_ = 0.28, F = 6.35, N = 15; Figure 4; qPCR threshold cycle for viral detection ranged from 14.3-23.8).

**Figure 3.**
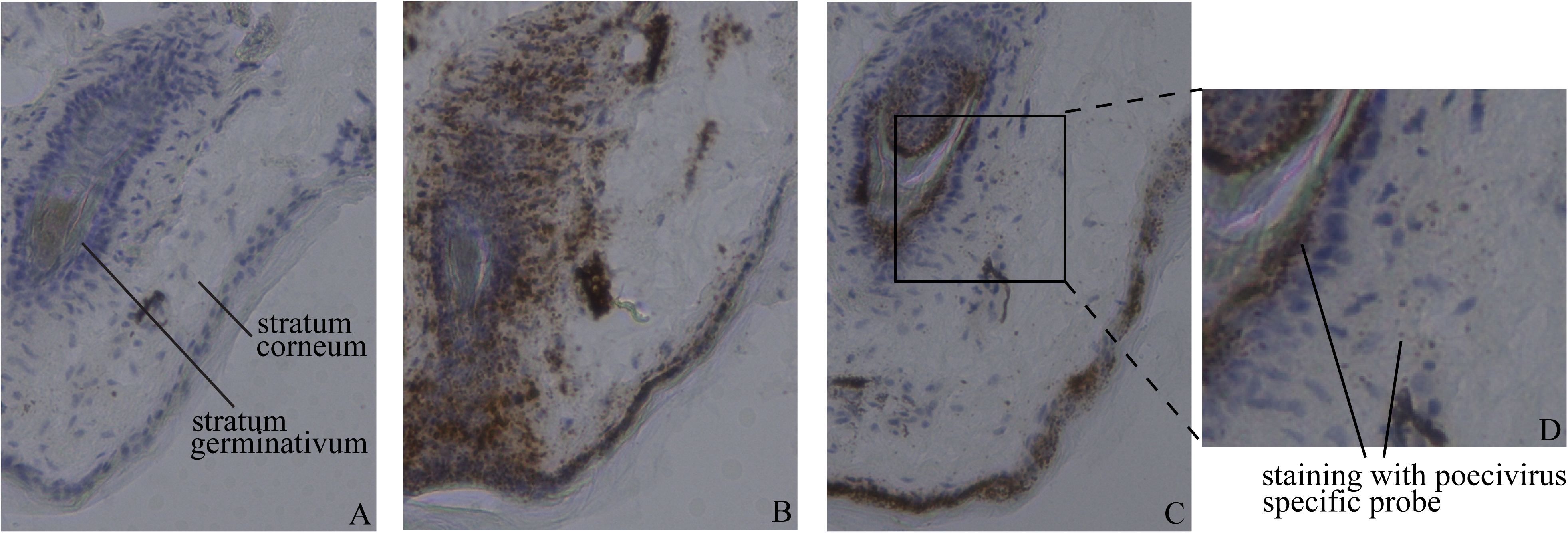
Viral localization. Transverse sections of beak of BCCH displaying AKD (beak length of 18.7 mm) and infected with Poecivirus, as determined by targeted PCR followed by Sanger sequencing. Cell nuclei are stained blue with hematoxylin and RNA sequence specific probes are stained brown. Panel A) section hybridized with probe for DapB bacterial gene (negative control); B) section hybridized with probe for BCCH NADH ND2 gene (positive control); C) section hybridized with probe for Poecivirus; D) inset of Panel C.

**Figure 4.**
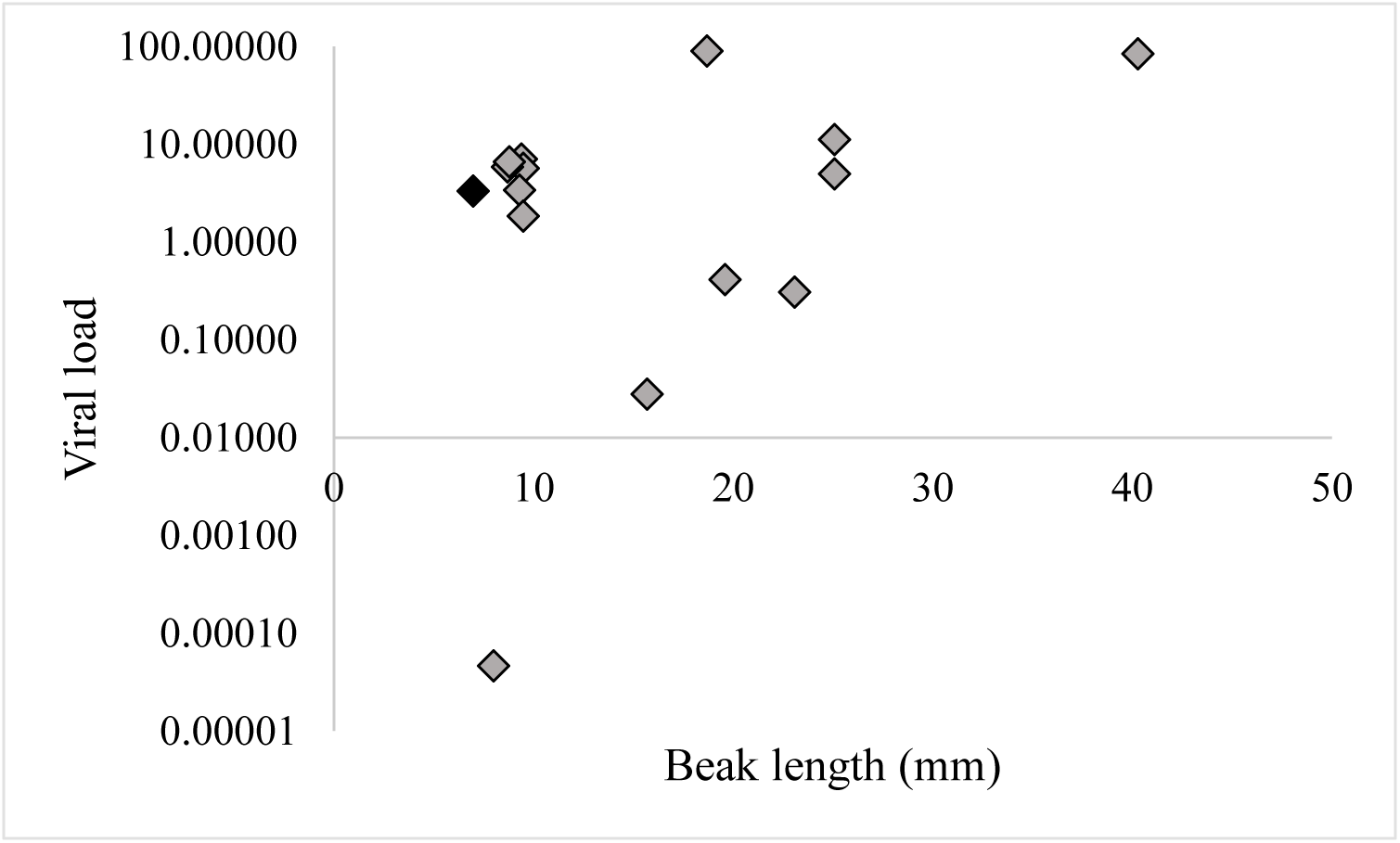
Viral load and beak length. Relative levels of viral RNA in beaks from Poecivirus-infected BCCH individuals were measured by qRT-PCR. Levels were normalized to levels of avian cellular RNA. Individuals with AKD are represented by grey diamonds, the single asymptomatic individual is shown in black.

We next sought to address the localization of Poecivirus at the site of pathology in AKD-affected individuals. Examination of beak tissue from four Poecivirus-infected individuals via RNAscope *in situ* hybridization revealed the presence of virus in the stratum germinativum and the stratum corneum (Figure 3). The same method failed to detect virus in the brain, liver, or gastrointestinal tract of one of these individuals in which we examined these tissues (data not shown).

As a positive strand virus, detection of Poecivirus negative strand is indicative of viral replication. To test for the presence of actively replicating virus, we carried out studies to directly detect and quantify both negative and positive sense Poecivirus RNAs. We used a total of 8 probes to detect negative sense virus RNA, 2 probes for positive sense virus RNA, and 2 probes for the BCCH housekeeping gene NADH ND2 (Figure 5). We detected expression of both positive and negative strand Poecivirus RNA in 11/11 Poecivirus-infected individuals (Figure 5). The probes for negative strand virus varied in their efficacy, detecting negative strand virus in 6-10 AKD individuals, although no probe to negative strand virus detected virus in every individual that tested positive for negative strand virus overall. One probe failed to detect virus in both the samples and the positive controls, including those that contained amounts of negative strand virus well above the stated minimum detection limits of the assay; as a result, this probe is absent from Figure 5. The number of negative strand RNAs normalized to the average number of BCCH NADH ND2 (when both probes to this gene were considered, GenBank: KF183899.1) ranged from 4.05E-04 to 1.13E-01. In comparison, the number of positive strand RNAs normalized to the average number of BCCH NADH ND2 was much higher, ranging from 1.09E-01 to 9.28E00. Neither negative nor positive strand RNAs explained a statistically significant amount of variation in beak length when normalized to average counts of the BCCH housekeeping gene (NADH ND2), (R^2^_adj_ = 0.013, and R^2^_adj_ <0.0001, respectively).

**Figure 5.**
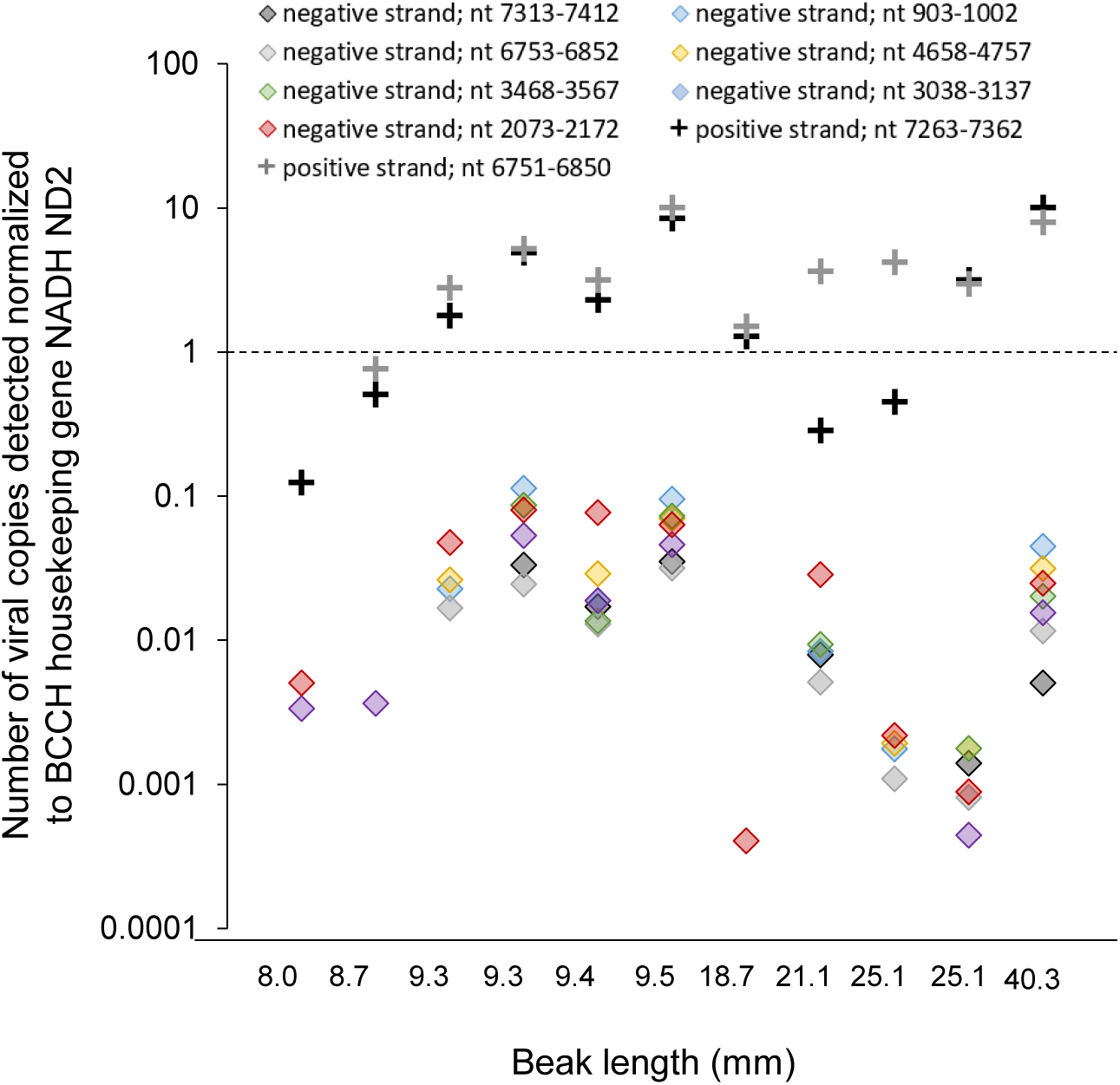
Viral replication. Relative levels of positive (pluses) and negative strand (diamonds) viral RNA in beaks from Poecivirus-infected BCCH (N = 11) were measured by a strand-specific gene expression assay. Levels were normalized to levels of avian cellular RNA. The presence of negative strand RNA indicates active viral replication. One probe to negative strand virus failed, and so does not appear in the graph. Probes are referred to by the position of the target nucleotides (nt) in the Poecivirus genome.

## DISCUSSION

In this study, we investigated the relationship between Poecivirus and avian keratin disorder (AKD), a disease characterized by beak deformities that appears to affect a large number of species across multiple continents. We previously used next-generation sequencing to identify and characterize the full-length genome of Poecivirus, a novel picornavirus, present in a small cohort of BCCH with AKD. Here we expand on that work, testing a substantially larger cohort of individuals (N = 124) for the presence of the virus. We demonstrated a statistically significant correlation between Poecivirus infection and AKD, with Poecivirus present in 29/29 (100%) of AKD-affected BCCH, and 9/95 (9.5%) of control individuals. These results are consistent with those of our previous study, which found an identical prevalence of Poecivirus in AKD-affected birds (100% of 29) and a slightly higher but not statistically different prevalence among asymptomatic control individuals (22% of 9; p = 0.29, χ^2^ = 1.14, df = 1). The slight discrepancy is likely attributable to the small sample size of control birds in our earlier study, although it could also reflect better detection of Poecivirus in sectioned tissue samples vs. swabs, or seasonal differences in disease spread or pathology, as has been documented in a variety of other infectious avian diseases (e.g., [15-20]).

The finding that some phenotypically AKD-asymptomatic (“control”) birds were positive for Poecivirus also merits further consideration. There are several potential explanations for this finding (for an in-depth discussion of this topic, see [7]). Briefly, the asymptomatic birds that tested positive for Poecivirus may have had subclinical infections (as has been suggested for related viruses [21]), or may have been in an early stage of infection, before sufficient time had passed for them to grow the elongated beak that defines AKD. Indeed, 1/4 (25%) individuals that tested positive for Poecivirus while apparently unaffected by AKD, and for which we have subsequent data, was later captured with an elongated beak. This individual was captured with a normal beak in April 2016 (beak length 7.4 mm) and tested positive for Poecivirus via cloacal swab; when recaptured 7 months later, it still tested positive for Poecivirus but now had a severely elongated beak (12.5 mm; we do not have data on the infection status of the other three birds upon recapture). Alternatively, it is possible that Poecivirus infection is not related to AKD; however, the presence of Poecivirus in significantly more AKD-affected birds than unaffected individuals suggests either that it is the causative agent of AKD or that AKD promotes or increases susceptibility to Poecivirus.

While we previously detected Poecivirus in beak tissue of BCCH with AKD, it was unknown whether Poecivirus was infecting cells of the avian beak; therefore, it remained a possibility that Poecivirus was not an avian virus but was simply present in beak samples because of incidental contact with the virus, for example, via foraging. The *in situ* hybridization data presented here indicate that Poecivirus was present in cells of the stratum corneum and stratum germinativum in four individuals with AKD. These data must be considered preliminary as we were unable to test control individuals for the presence of Poecivirus in beak tissue; however, the presence of this virus in avian cells provides further evidence that Poecivirus is indeed an infectious avian virus rather than an incidental contaminant. This evidence that Poecivirus is an avian virus is consistent with phylogenetic analysis, which shows that Poecivirus’ closest relatives are the avian megriviruses [7]. In addition, the presence of Poecivirus in beak tissue, but not other tissues (brain, liver, gastrointestinal tract), is in keeping with the focal pathology of AKD, which primarily affects beak tissue. Moreover, the localization of Poecivirus in the stratum germinativum, the layer of the beak that gives rise to keratin cells, is consistent with the pathology of AKD, which is characterized by overgrowth of the keratin layer of the beak. Our previous study detected low levels of virus in brain, liver, and gastrointestinal tract; however, it is possible that the virus was present in these tissues incidentally and was not actively replicating.

The gene expression data presented here indicate that Poecivirus is actively replicating in avian beak tissue. Poecivirus, like other picornaviruses, is a nonenveloped single stranded positive sense RNA virus. As with other picornaviruses, the Poecivirus genome encodes for an RNA-dependent RNA polymerase that makes complementary minus strands of RNA during viral replication [22]. These negative strand RNAs are only present during viral replication. Therefore, the presence of negative strand Poecivirus RNA in the beak tissue of all AKD-affected individuals tested (11/11) strongly suggests that Poecivirus was actively replicating in these individuals.

Our data suggest that field studies of Poecivirus and AKD would benefit by combining both cloacal and buccal swabs to test for the presence of Poecivirus in wild avian populations (Table 1). Cloacal swabs performed best in non-terminal field-based sampling of AKD individuals for the presence of Poecivirus; cloacal swabs detected Poecivirus in 28/29 (96.6%) of AKD birds. Meanwhile, buccal swabs appear to be prone to type II error (false negatives) in individuals with AKD, detecting Poecivirus in only 3/21 (14.3%) AKD individuals tested; however, one of these individuals tested negative for the presence of Poecivirus via cloacal swab, indicating that combining the two methods could provide additional information. In contrast, 4/9 (44%) control individuals that tested positive for Poecivirus did so via buccal swab only. The difference in efficacy of the two swab types for detecting Poecivirus between AKD and control individuals is intriguing; future studies should investigate whether this difference has a biological basis. For example, detection of Poecivirus in cloacal swabs may require viral shedding, which could increase later in infection, once the elongated beak has had time to develop. While we tested a relatively small number of blood and fecal samples for the presence of Poecivirus, neither of these sample types showed promise for Poecivirus detection; this could be the result of PCR inhibitors known to be present in both feces and blood [23].

## CONCLUSIONS

Poecivirus continues to warrant further investigation as a candidate agent of AKD. The data presented here show a strong, statistically significant relationship between Poecivirus infection and AKD, and provide evidence that Poecivirus is indeed an avian virus, infecting and actively replicating in beak tissue of AKD-affected BCCH. Cloacal and buccal swabs provide a promising method for non-terminal field-testing of wild birds for the presence of Poecivirus, and a way forward for future studies of the impacts of this pathogen on the ecology and fitness of wild bird populations. Ultimately, a viral challenge of healthy individuals with Poecivirus is needed to determine with certainty the role of Poecivirus in AKD and will allow us to better understand the impact of this disease on avian populations worldwide.

## LIST OF ABBREVIATIONS

AKD: avian keratin disorder
BCCH: black-capped chickadee (*Poecile atricapillus*)

## DECLARATIONS

Ethics approval and consent to participate: All work was conducted with the approval of the U. S. Geological Survey (USGS) Alaska Science Center Institutional Animal Care and Use Committee (Assurance #2016-14) and under appropriate state and federal permits.

### Consent for publication

Not applicable

### Availability of data and material

The datasets generated and/or analyzed during the current study are available in [24].

### Funding

A National Science Foundation Postdoctoral Research Fellowship (M.Z.)

The Howard Hughes Medical Institute (J.L.D.)

U. S. Geological Survey through the Wildlife Program of the Ecosystems Mission Area (C.M.H., C.V.)

The Chan Zuckerberg Biohub (J.L.D.)

The funders had no role in study design, data collection and interpretation, or the decision to submit the work for publication. Any use of trade, product, or firm names in this publication is for descriptive purposes only and does not imply endorsement by the U. S. Government.

### Authors’ contributions

M.Z. participated in the conception, design, and coordination of the study, performed lab work and data analyses, and wrote and revised the manuscript; C.V. participated in the conception, design, and coordination of the study, collected the samples, and participated in revising the manuscript; C.M.H. and J.D. oversaw the conception and design of the project, supervised its execution and participated in revising the manuscript.

## Acknowledgments

We thank Hannah Retallack, Christopher Liverman, and Caleigh Mandel-Brehm for laboratory assistance; Kristeene Knopp, Amy Kistler, and Andrew Reeves for comments on an earlier version of this manuscript; Jennifer Mann for logistical assistance; and Lisa Pajot, Rachel Richardson, and numerous volunteers for assistance with sample collections.

## Additional files

File name: Additional_file_1.pdf

File format: .pdf

Title of data: PCR primers

Description of data: PCR primers used to detect Poecivirus

## References

1. Handel CM, Pajot LM, Matsuoka SM, Hemert CV, Terenzi J, Talbot SL, Mulcahy DM, Meteyer CU, Trust KA: Epizootic of beak deformities among wild birds in Alaska: An emerging disease in North America? Auk 2010, 127:882–898.

2. Van Hemert C, Handel CM, O’Hara TM: Evidence of accelerated beak growth associated with avian keratin disorder in black-capped chickadees (Poecile atricapillus). Journal of Wildlife Diseases 2012, 48:686–694.

3. Van Hemert C, Armién AG, Blake JE, Handel CM, O’Hara TM: Macroscopic, histologic, and ultrastructural lesions associated with avian keratin disorder in black-capped chickadees (Poecile atricapillus). Veterinary Pathology Online 2013.

4. D’Alba L, Van Hemert C, Spencer KA, Heidinger BJ, Gill L, Evans NP, Monaghan P, Handel CM, Shawkey MD: Melanin-based color of plumage: Role of condition and of feathers’ microstructure. Integrative and Comparative Biology 2014, 54:633–644.

5. Wilkinson LC, Handel CM, Van Hemert C, Loiseau C, Sehgal RNM: Avian malaria in a boreal resident species: Long-term temporal variability, and increased prevalence in birds with avian keratin disorder. International Journal for Parasitology 2016, 46:281–290.

6. Van Hemert C, Handel C: Beak deformities in northwestern crows: evidence of a multispecies epizootic. Auk 2010, 127:746–751.

7. Zylberberg M, Van Hemert C, Dumbacher JP, Handel CM, Tihan T, DeRisi JL: Novel picornavirus associated with avian keratin disorder in Alaskan birds. mBio 2016, 7.

8. Big Garden Beak Watch Results [http://www.bto.org/volunteer-surveys/gbw/about/background/projects/bgbw/results/species]

9. Harrison T: Beak deformities of garden birds. British Birds 2011, 104:538–541.

10. Craves J: Passerines with deformed bills. North American Bird Bander 1994, 19:14–18.

11. Tully T, Jr, Lawton M, Dorrestein G: Handbook of Avian Medicine. 2nd edition edn. New York, NY: Butterworth-Heinemann, Elsevier Science Limited; 2000.

12. Handel CM, Van Hemert C: Environmental contaminants and chromosomal damage associated with beak deformities in a resident North American passerine. Environ Toxicol Chem 2015, 34:314–327.

13. Longmire JL, Lewis AK, Brown NC, Buckingham JM, Clark LM, Jones MD, Meincke LJ, Meyne J, Ratliff RL, Ray FA, et al Isolation and molecular characterization of a highly polymorphic centromeric tandem repeat in the family falconidae. Genomics 1988, 2:14–24.

14. Gibson DG, Young L, Chuang R-Y, Venter JC, Hutchison CA, Smith HO: Enzymatic assembly of DNA molecules up to several hundred kilobases. Nat Meth 2009, 6:343–345.

15. Altizer S, Dobson A, Hosseini P, Hudson P, Pascual M, Rohani P: Seasonality and the dynamics of infectious diseases. Ecology Letters 2006, 9:467–484.

16. Cornelius J, Zylberberg M, Breuner C, Gleiss AC, Hahn T: Assessing the role of reproduction and stress in the spring emergence of Haematozoan parasites in birds. The Journal of Experimental Biology 2014, 217:841–849.

17. Zylberberg M, Lee KA, Klasing KC, Wikelski M: Increasing avian pox prevalence varies by species, and with immune function, in Galápagos finches. Biological Conservation 2012, 153:72–79.

18. Zylberberg M, Lee KA, Klasing KC, Wikelski M: Variation with land use of immune function and prevalence of avian pox in Galapagos finches. Conservation Biology 2013, 27:103–112.

19. Wang R-H, Jin Z, Liu Q-X, van de Koppel J, Alonso D: A simple stochastic model with environmental transmission explains multi-year periodicity in outbreaks of avian flu. PLoS ONE 2012, 7:e28873.

20. Jahangir A, Ruenphet S, Ueda S, Ueno Y, Shoham D, Shindo J, Okamura M, Nakamura M, Takehara K: Avian influenza and Newcastle disease viruses from northern pintail in Japan: Isolation, characterization and inter-annual comparisons during 2006–2008. Virus Research 2009, 143:44–52.

21. Boros Á, Nemes C, Pankovics P, Kapusinszky B, Delwart E, Reuter G: Identification and complete genome characterization of a novel picornavirus in turkey (Meleagris gallopavo). Journal of General Virology 2012, 93:2171–2182.

22. Ehrenfeld E, Domingo E, Roos RP: The Picornaviruses. Washington, DC: American Society for Microbiology Press; 2010.

23. Rådström P, Knutsson R, Wolffs P, Lövenklev M, Löfström C: Pre-PCR processing. Molecular Biotechnology 2004, 26:133–146.

24. Zylberberg M, C. Van Hemert, C.M. Handel, and J.L. DeRisi.: Genetic data associated with avian keratin disorder and Poecivirus in black-capped chickadees, Alaska, 2016-2017. (release USGS ed.; 2018.

