## Supplementary Materials for "Avian keratin disorder of Alaska black-capped chickadees is associated with Poecivirus infection"

| Primer pair | Primer name | Sequence |
|---|---|---|
| 1 | Poeci_1F | TGGCTGCTCTAGAGGATAAAGG |
| | Poeci_1R | ACTGCACTACAACCAAATCTGT |
| 2 | Poeci_2F | AGCTTGGCCCCTCTAATTGT |
| | Poeci_2R | GATTACTGTTCCGGTCTCTTGG |
| 3 | Poeci_3F | TGTCATACTTGCCACCTCCG |
| | Poeci_3R | GCAACTTCCAATTGCACGTC |
| 4 | Poeci_4F | TGGGCATTGTCTCGAGTGTA |
| | Poeci_4R | TACGAAAAGCCTCAGTCGGA |
| 5 | Poeci_5F | GACCGTGGATAATTATGTGAAAGGATTGAGACGT |
| | Poeci_5R | GCGAACAGTGGTAGATACAGGCCGC |
| 6 | Poeci_6F | AAGCTCCATATGATCCAAATTATTCGCGGCG |
| | Poeci_6R | AAGCAATATTATTCACCTCAATCAACTGTACCACA |
| 7 | Poeci_7F | CAAAGTGTTGTAGAGGCGGC |
| | Poeci_7R | ACAACAAAATCCCGCAACGT |
| 8 | avi_8F | CAAGCTGCACCAAGGGAAAT |
| | avi_8r | TCACAGAATCACTAGGTTGGAAG |

Additional file 1: primers. Poeci_2F targeting the 5’ UTR is not represented in the Sanger sequencing validated Poecivirus genome deposited in GenBank (accession number KU977108).
